# Evolution of antibody cross-reactivity to influenza H5N1 neuraminidase from an N2-specific germline

**DOI:** 10.1101/2025.06.20.660733

**Authors:** Huibin Lv, Yang Wei Huan, Qi Wen Teo, Chunke Chen, Tossapol Pholcharee, Akshita B. Gopal, Madison R. Ardagh, Jessica J. Huang, Ruipeng Lei, Xin Chen, Yuanxin Sun, Yun Sang Tang, Arjun Mehta, Mateusz Szlembarski, Kevin J. Mao, Emily X. Ma, Lucas E. Wittenborn, Meixuan Tong, Lucia A. Rodriguez, Letianchu Wang, Chris K. P. Mok, Nicholas C. Wu

## Abstract

The ongoing spread of highly pathogenic avian influenza H5N1 clade 2.3.4.4b virus in animals and its occasional spillover to humans have raised concerns about a potential H5N1 pandemic. Although recent studies have shown that pre-existing human antibodies can recognize H5N1 neuraminidase, the molecular basis of how this cross-reactivity develops remains poorly understood. In this study, we used a phage display antibody library derived from 245 healthy donors to isolate an antibody, HB420, that cross-reacts with neuraminidases of human H3N2 and avian H5N1 clade 2.3.4.4b viruses and confers protection *in vivo*. Cryo-EM analysis reveals that HB420 targets the neuraminidase active site by mimicking sialic acid binding through a single Asp residue. Furthermore, the inferred germline of HB420 is N2-specific but acquires cross-reactivity to H5N1 neuraminidase through somatic hypermutation. Overall, our findings provide insights into how neuraminidase antibody evolves breadth, which has important implications for the development of broadly protective influenza vaccines.

## INTRODUCTION

Influenza A viruses remain a major public health and socioeconomic threat. Influenza epidemics are responsible for around 3 to 5 million cases of severe illness and 290,000 to 650,000 deaths worldwide each year.^1^ During pandemic years, influenza A viruses can even cause up to millions of deaths globally.^2^ Influenza A viruses have two major surface antigens, hemagglutinin (HA) and neuraminidase (NA). HA mediates host cell entry by binding to sialylated receptors and mediating virus-host membrane fusion, whereas NA facilitates viral release by cleaving sialic acids. HA and NA are classified into 19 (H1-H19) and 11 (N1-N11) subtypes, respectively. While H17-H18 and N10-N11 have only been observed in bats,^3,4^ all other subtypes are found in aquatic birds, which represent the natural reservoir of influenza A viruses.^5,6^ Currently, H1N1 and H3N2 subtypes are circulating in human. Nevertheless, other avian influenza subtypes have occasionally spilled over to humans.^7^ In 2024, the widespread detection of highly pathogenic avian influenza (HPAI) H5N1 clade 2.3.4.4b virus in the US dairy cattle^8–11^ raised concerns about a potential H5N1 pandemic.^12^ The threat has been further intensified by multiple cases of HPAI H5N1 human infection in the US, including one fatality.^13,14^

Seasonal influenza vaccines, which are currently the most effective preventive measure against influenza virus infection, do not confer protection against zoonotic influenza subtypes.^15^ Therefore, influenza vaccines that provide broad protection against multiple human and zoonotic subtypes are urgently needed for pandemic preparedness. Due to the discovery of broadly neutralizing human antibodies targeting the HA nearly two decades ago,^16^ the development of broadly protective influenza vaccines has largely focused on HA. Nevertheless, several broadly protective human antibodies targeting neuraminidase (NA) have also been reported in recent years, including some that cross-react with both group 1 (N1, N4, N5, and N8) and group 2 (N2, N3, N6, N7, and N9) NAs,^17^ as exemplified by FNI9,^18^ 4N2C402,^19^ 1G01,^20^ Z2B3,^21^ DA03E17,^22^ NCS1.2,^23^ and mAb-297.^24^ The discovery of broadly protective NA antibodies demonstrates the importance of NA as a target for the development of broadly protective influenza vaccines.

Multiple studies suggest that NA antibody responses, albeit poorly elicited by seasonal influenza vaccines,^25^ are critical for protection against zoonotic subtypes. For example, pre-existing antibodies to the H2N2 NA have been shown to confer protection during the 1968 H3N2 pandemic.^26^ More recently, serology studies have demonstrated that while humans have minimal pre-existing HA-neutralizing antibodies against H5N1 clade 2.3.4.4b virus, pre-existing NA-inhibiting antibodies against the virus are prevalent.^27,28^ These findings suggest that pre-existing NA antibodies are likely to play a key role in reducing morbidity and mortality in the event of an H5N1 pandemic. However, the molecular basis for the development of pre-existing NA antibodies against H5N1 clade 2.3.4.4b virus remains largely elusive.

In this study, we constructed a phage display antibody library using the peripheral blood mononuclear cells (PBMCs) from 245 healthy donors and screened it against H5N1 NA. This effort led to the identification of one cross-group antibody, HB420, that not only binds to H5N1 NA but also to H3N2 NA. While HB420 exhibited much stronger NA inhibition activity against H3N2 NA than H5N1 NA and inhibited the growth of H3N2 virus but not H5N1 virus *in vitro*, it protected against both H3N2 virus and H5N1 virus *in vivo*. Structural analysis by cryo-electron microscopy (cryo-EM) further revealed that HB420 targeted the conserved NA active site through receptor mimicry, mediated by a single Asp residue in the complementarity-determining region (CDR) H3. While the inferred germline of HB420 could not bind to H5N1 NA, its binding to H3N2 NA remained strong. Our mutagenesis results indicate that HB420 developed its cross-reactivity against H5N1 NA through somatic hypermutations (SHMs) in the heavy chain. Throughout this study, N2 numbering and Kabat numbering are used for NA residue positions and antibody residue positions, respectively, unless otherwise stated.

## RESULTS

### Pre-existing NA antibodies in humans against zoonotic influenza A subtypes

To assess pre-existing NA antibodies against zoonotic influenza A subtypes, we analyzed plasma samples from 245 healthy donors collected in Hong Kong between January and March 2020. Donors were stratified into five age groups: 16-20, 21-30, 31-40, 41-50, and 51-67 years. The binding activity of these plasma samples to recombinant NAs were evaluated by enzyme-linked immunosorbent assay (ELISA). This experiment included the NAs from A/cattle/Texas/56283/2024 (Tx24, H5N1), A/Jiangsu/428/2021 (Js21, H10N3), A/Jiangsu/1/2018 (Js18, H7N4), and A/mallard/Alberta/362/2017 (Alb17, H3N8). Additionally, NAs from A/Michigan/45/2015 (Mich15, H1N1) and A/Singapore/INFIMH-16-0019/2016 (Sing16, H3N2), which are the seasonal human strains, were included as positive NA controls. We also included five plasma samples from one-year-old infants with no history of influenza infection or vaccination^29^ as baseline controls to establish the cutoff for binding detection.

As expected, most individuals harbored detectable antibodies against Mich15 (H1N1) NA and Sing16 (H3N2) NA. Notably, antibody levels against both seasonal NAs were significantly higher in the 16-20 age group compared to the 51-67 age group, with *p* < 0.05 for Mich15 (H1N1) NA and *p* < 0.0001 for Sing16 (H3N2) NA (**Figure 1A-B**). Consistent with previous studies,^27,28,30,31^ antibodies against Tx24 (H5N1) NA were also detected, with younger age groups exhibiting higher antibody levels compared to older age groups (**Figure 1C**). By contrast, antibody levels against Js21 (H10N3) NA, Js18 (H7N4) NA, and Alb17 (H3N8) NA were comparable across all age groups. Although several plasma samples contained antibodies against Js21 (H10N3) NA (**Figure 1D**), most plasma samples had no antibody binding to Js18 (H7N4) NA, and Alb17 (H3N8) NA (**Figure 1D-1F**). These findings substantiate that humans have pre-existing NA antibodies against some zoonotic influenza A subtypes, which were likely elicited by seasonal influenza virus infection.^27,28,30,31^

**Figure 1.**
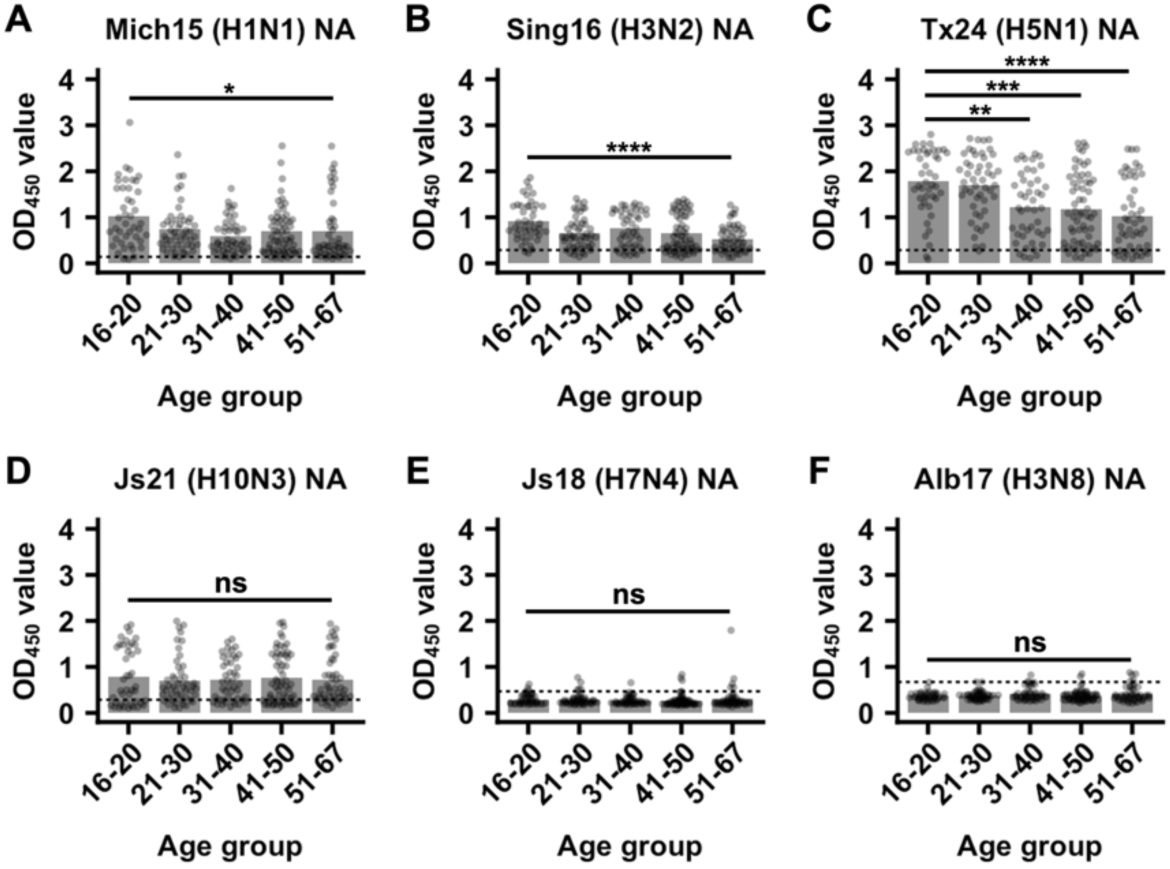
Serology analysis of different influenza NA subtypes in healthy donors. Antibody binding activity of plasma samples against NAs from six different influenza virus strains, including **(A)** A/Michigan/45/2015 (H1N1), **(B)** A/Singapore/INFIMH-16-0019/2016 (H3N2), **(C)** A/cattle/Texas/56283/2024 (H5N1), **(D)** A/Jiangsu/428/2021 (H10N3), **(E)** A/Jiangsu/1/2018 (H7N4), and **(F)** A/Mallard/Alberta/362/2017 (H3N8) were measured by ELISA. Five plasma samples from 1-year-old infants without any history of influenza infection^29^ were used as negative controls. The dashed lines indicate the cutoff for positive binding, which is defined as twice the OD_450_ value of the negative control. P-values were calculated using an unpaired two-tailed Students t-test (*P < 0.05, **P < 0.01, ***P < 0.001, ****P < 0.0001; ns, not significant).

### Isolating a monoclonal antibody against H5N1 NA by phage display

To understand the molecular basis of how humans developed pre-existing NA antibodies against Tx24 (H5N1) NA, we constructed a phage display antibody library using PBMCs from the same 245 healthy donors used in our serology analysis (**Figure 1**). After two rounds of phage panning of the antibody library against Tx24 (H5N1) NA, 20 clones were sequenced. All 20 clones represented the same antibody, which was encoded by *IGHV1-69*/*IGLV2-14*. We designated this antibody as HB420 (**Figure 2A**).

**Figure 2.**
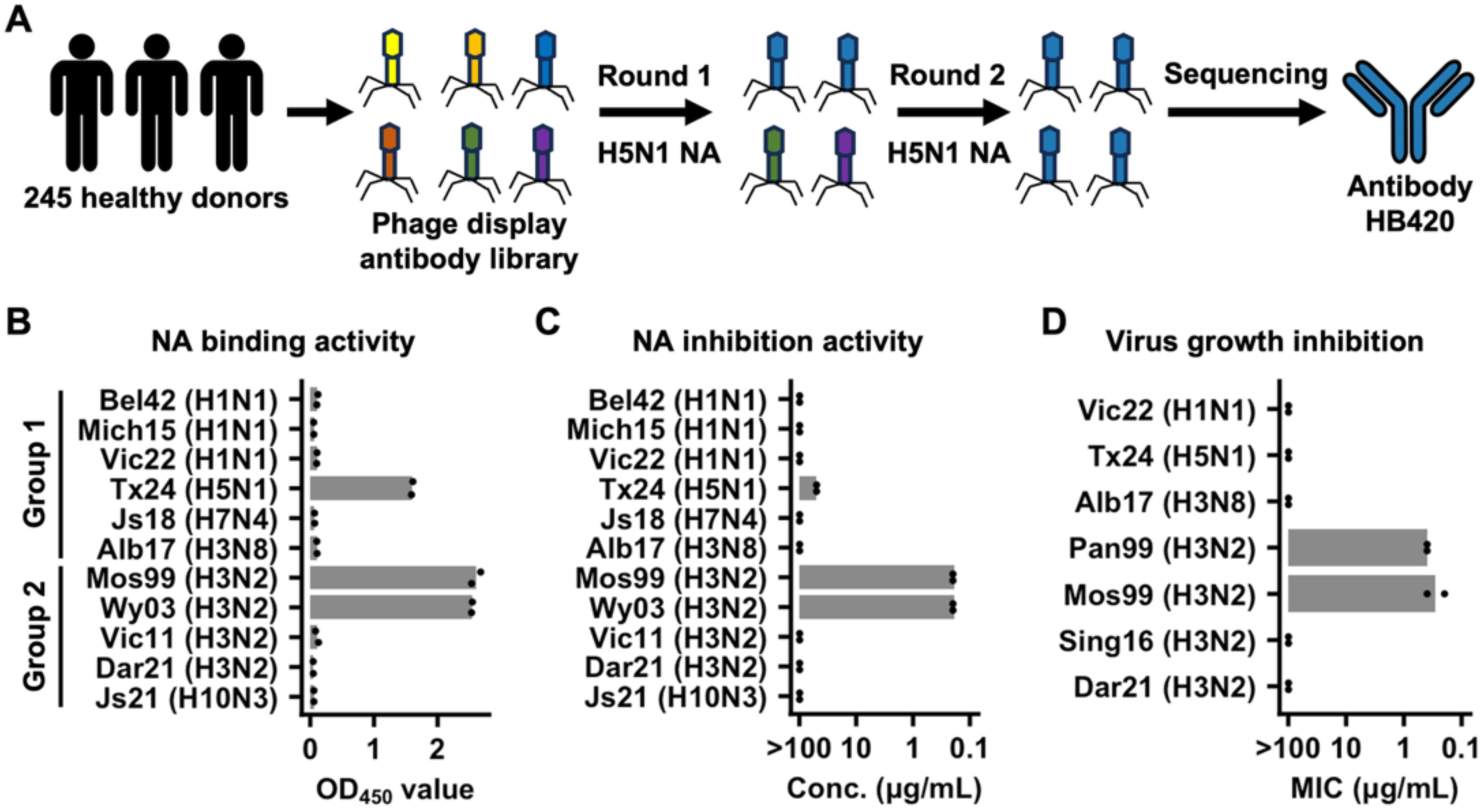
Discovery and *in vitro* characterization of HB420. **(A)** Schematic workflow for identifying HB420 from a phage display library derived from 245 healthy donors. **(B)** Binding activity of HB420 against recombinant NAs from the indicated strains was measured by ELISA. **(C)** NA inhibition activity of HB420 against recombinant NAs from the indicated strains was measured by ELLA. **(D)** Virus growth inhibition activity of HB420 against the indicated strains was measured. Minimum inhibitory concentration (MIC) is shown. **(B-D)** Each bar represents the mean of two technical replicates. Each datapoint represents one technical replicate. Data from one of two experiments with similar results are shown. Bel42: A/Bel/1942 (H1N1), Mich15: A/Michigan/45/2015 (H1N1), Vic22: A/Victoria/4897/2022 (H1N1), Tx24: A/cattle/Texas/56283/2024 (H5N1), Js18: A/Jiangsu/1/2018 (H7N4), Alb17: A/mallard/Alberta/362/2017 (H3N8), Mos99: A/Moscow/10/1999 (H3N2), Wy03: A/Wyoming/3/2003 (H3N2), Vic11: A/Victoria/361/2011 (H3N2), Dar21: A/Darwin/6/2021 (H3N2), and Js21: A/Jiangsu/428/2021 (H10N3). All recombinant viruses were 7:1 reassortants, containing the NA segment from the indicated strains on the backbone of A/Puerto Rico/8/1934 (PR8, H1N1).

To probe the cross-reactivity of HB420, its binding activity was tested against a panel of NAs from different subtypes using ELISA (**Figure 2B**). Aside from Tx24 (H5N1) NA, HB420 did not bind to any other group 1 NAs tested, including those from N1 subtype. Yet, HB420 bound to a subset of N2 NAs, including A/Moscow/10/1999 (Mos99, H3N2) and A/Wyoming/03/2003 (Wy03, H3N2). The binding activity of HB420 to Mos99 (H3N2) NA and Wy03 (H3N2) NA was even stronger than to Tx24 (H5N1) NA. We further measured the ability of HB420 to inhibit NA activity using an enzyme-linked lectin assay (ELLA) (**Figure 2C**). Consistent with its binding activity, HB420 had strong inhibitory activity against both Mos99 (H3N2) NA and Wy03 (H3N2) NA, with a 50% inhibitory concentration (IC_50_) of 0.19 μg/mL. HB420 also had weak inhibitory activity against Tx24 (H5N1) NA, with an IC_50_ of 50 μg/mL. Additionally, HB420 inhibited the growth of Mos99 (H3N2) virus and A/Panama/2007/1999 (Pan99, H3N2) virus, with minimum inhibitory concentrations (MICs) of 0.19 and 0.39 μg/mL, respectively. By contrast, even at 100 μg/mL, HB420 did not inhibit the growth of other influenza A viruses tested, including one containing Tx24 (H5N1) NA (**Figure 2D**). These results demonstrated that HB420 was a cross-group antibody, despite having stronger functional activity against N2 NA.

### HB420 confers protection *in vivo* against both N1 and N2 viruses

Given the *in vitro* cross-reactivity of HB420, we next evaluated its prophylactic protective efficacy *in vivo* against 7:1 reassortant influenza viruses bearing either Mos99 (H3N2) NA or Tx24 (H5N1) NA on the PR8 (H1N1) backbone. Briefly, HB420 was administered to female BALB/c mice via intraperitoneal injection either 4 hours prior to (prophylactic) or 24 hours after (therapeutic) intranasal infection with 5×LD_50_ of virus. At 5 mg/kg, HB420 conferred complete protection against virus bearing the Mos99 (H3N2) NA, with 100% survival and minimal weight loss in both prophylactic and therapeutic setting (**Figure 3A-B**). Consistently, lung viral titers measured on day 3 post-infection were reduced by four logs and two logs in HB420 prophylactically treated and therapeutically treated mice, respectively, compared to mice without antibody treatment (**Figure 3C**). At 10 mg/kg, HB420 also provided partial prophylactic protection against the recombinant virus bearing the Tx24 (H5N1) NA, with a survival rate of 40% (**Figure 3D-E**) and a two-log reduction in lung viral titers on day 3 post-infection (**Figure 3F**). Our results substantiated the notion that NA antibodies with no detectable virus growth inhibition activity *in vitro* could still be protective *in vivo*.^32–35^

**Figure 3.**
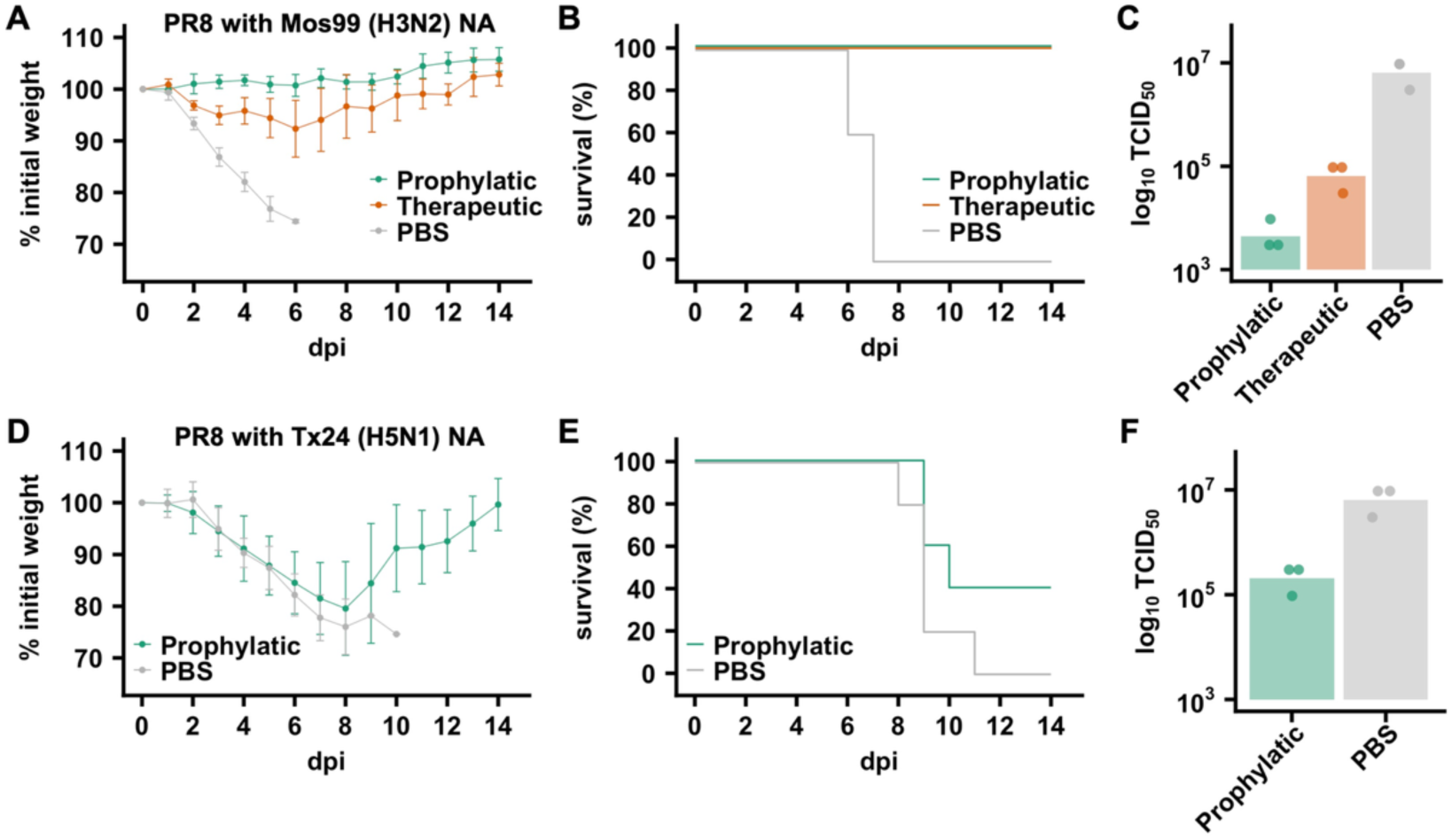
*In vivo* protection activity of HB420. HB420 at **(A-C)** 5 mg/kg or **(D-F)** 10 mg/kg was administered intraperitoneally to 6-week-old BALB/c mice either 4 hours prior to (prophylactic) or 24 hours after (therapeutic) challenge with 5×LD_50_ of recombinant A/Puerto Rico/8/1934 (H1N1) virus carrying **(A-C)** the NA from A/Moscow/10/1999 (H3N2) or **(D-F)** A/cattle/Texas/56283/2024 (H5N1). Representative experiments from two independent replicates with similar results are shown. **(A, D)** The mean percentage of body weight change post-infection is shown (n = 5). The humane endpoint was defined as a weight loss of 25% from initial weight on day 0. **(B, E)** Kaplan-Meier survival curves are shown (n = 5). **(C)** Lung viral titers on day 3 post-infection are shown (n = 3). Each bar represents the mean.

### HB420 targets the conserved NA active site

To elucidate the binding mode of HB420 to NA, we determined the cryo-EM structure of Mos99 (H3N2) NA in complex with the HB420 fragment antigen-binding (Fab) at a resolution of 2.16 Å (**Figure 4A and Table S1**). HB420 inserted its complementarity-determining region (CDR) H3 into the active site of NA (**Figure 4B**), with its epitope extending to the periphery of the active site (**Figure 4C**). HB420 heavy chain had a higher contribution to the paratope (841 Å^2^) compared to its light chain (427 Å^2^), with the CDR H3 alone accounting for 60% (510 Å^2^) of the heavy chain paratope (**Figure S1**). The CDR H3 of HB420 interacted with the NA active site mainly via electrostatic interactions and H-bonds (**Figure 4D**). At the tip of the CDR H3, the side chain of V_H_ D100a electrostatically interacted with NA R118, R292, and R371. Additionally, the backbone amide and carbonyl groups of V_H_ D100a H-bonded with NA D151 and H347, respectively. The CDR H3 of HB420 also H-bonded with NA R152, R430, and K431. Besides, V_H_ Y100d formed π-π stacking interaction with NA H347. The interactions between other CDRs of HB420 and NA were much less extensive, despite the presence of several H-bonds involving CDR H1, H2, L2, and L3 (**Figure 4E-4H**). As a result, our structural analysis demonstrated that HB420 inhibited the enzymatic activity of NA by directly blocking its active site.

**Figure 4.**
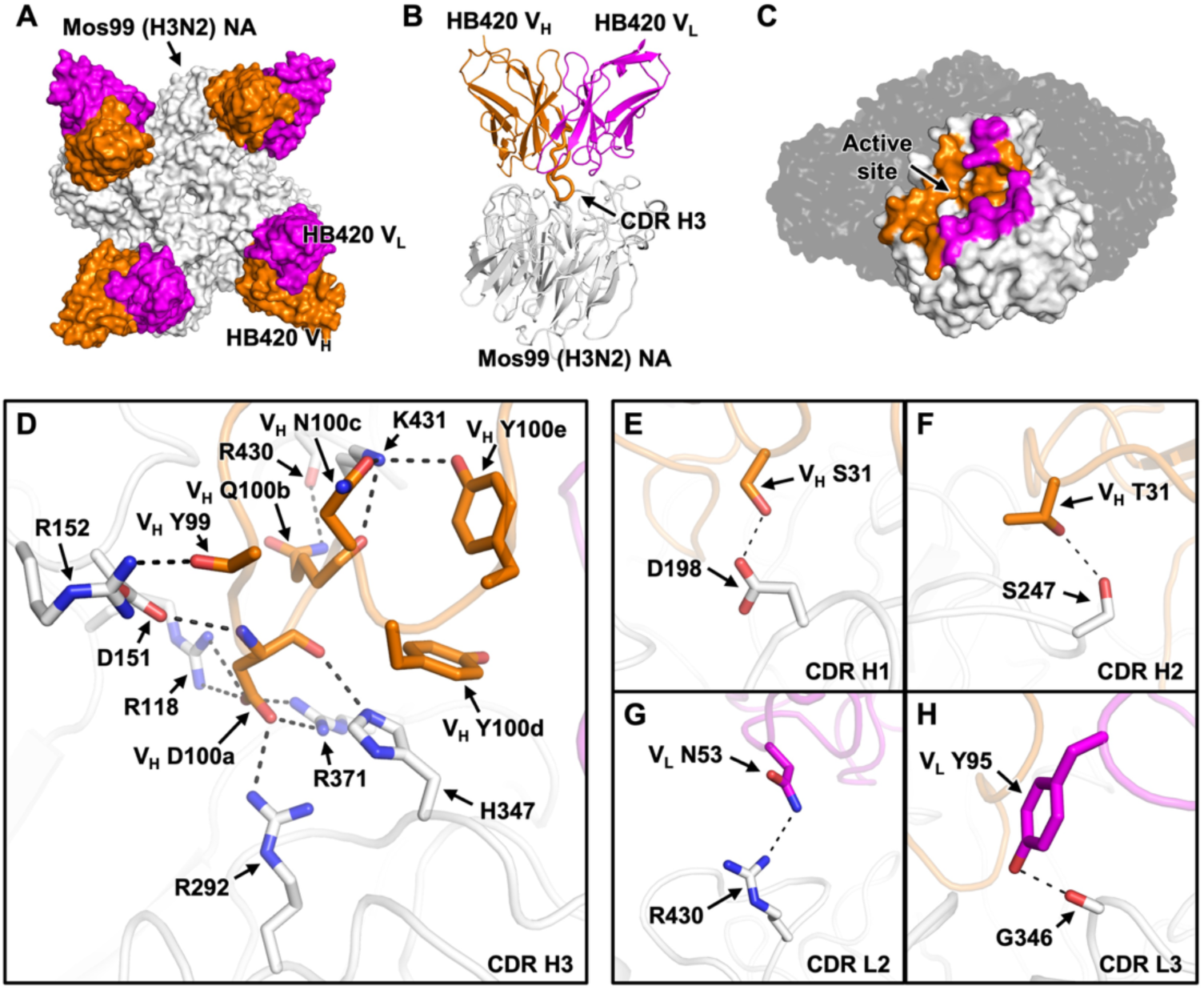
Structural analysis of HB420. **(A)** Cryo-EM structure of HB420 in complex with NA from A/Moscow/10/1999 (H3N2). HB420 and NA are shown in surface representation. NA is in white. Heavy chain variable domain (V_H_) and light chain variable domain (V_L_) are in orange and magenta, respectively. **(B)** HB420 and NA are shown in cartoon representation. CDR H3 of HB420 is indicated. Only one protomer is shown. **(C)** HB420 epitope on one protomer of NA is colored with V_H_ contacts in orange and V_L_ contacts in magenta. NA active site is indicated. **(D-H)** Interactions of NA with **(D)** CDR H3, **(E)** CDR H1, **(F)** CDR H2, **(G)** CDR L1, and **(H)** CDR L3 are shown. H-bonds and electrostatic interactions are represented by black dashed lines.

Certain key epitope residues of HB420 exhibited sequence variation across different influenza A virus strains (**Figure S2A**), which could explain the lack of binding of HB420 to some of the tested NAs (**Figure 2B**). For example, the π-π stacking interaction between V_H_ Y100d of HB420 and H347 of Mos99 (H3N2) NA would be abolished in human H1N1 NAs, which possessed an Asn at residue 347 instead (**Figure S2A-B**). This sequence variation at residue 347 was consistent with the lack of binding of HB420 to the NAs from more recent human H1N1 strains (**Figure 2B**). Moreover, the electrostatic interaction between V_H_ D55 of HB420 and K221 of Mos99 (H3N2) NA would become repulsive in NAs from more recent human H3N2 strains, which possessed acidic amino acids at residue 221 instead (**Figure S2A and S2C**). Since 2016, human H3N2 NA has acquired an N-glycosylation site at residue 245.^36^ Our structural analysis indicated that introducing an N-glycosylation site at residue 245 would prevent binding of HB420 due to steric hindrance (**Figure S2D**). These sequence variations at residues 221 and 245 were consistent with the lack of binding of HB420 to the NAs from recent human H3N2 strains (**Figure 2B**).

### Receptor mimicry by a single Asp residue in HB420

The structures of several cross-group human antibodies targeting the NA active site have been previously determined, including Z2B3,^37^ mAb-297,^24^ FNI9,^18^ 1G01,^38^ and 4N2C402.^19^ Same as HB420, Z2B3, mAb-297, and FNI9 are encoded by *IGHV1-69* (**Figure 5A**). Z2B3 is even encoded by the same light-chain germline gene as HB420. Nevertheless, these *IGHV1-69* antibodies had very different angles of approach to NA (**Figure 5B and Figure S3**). Relatedly, the CDR H3 conformations of these NA antibodies were very different (**Figure 5C**). These observations suggested that the binding modes of antibodies to NA active site were mainly determined by the CDR H3 rather than the germline gene usage.

**Figure 5.**
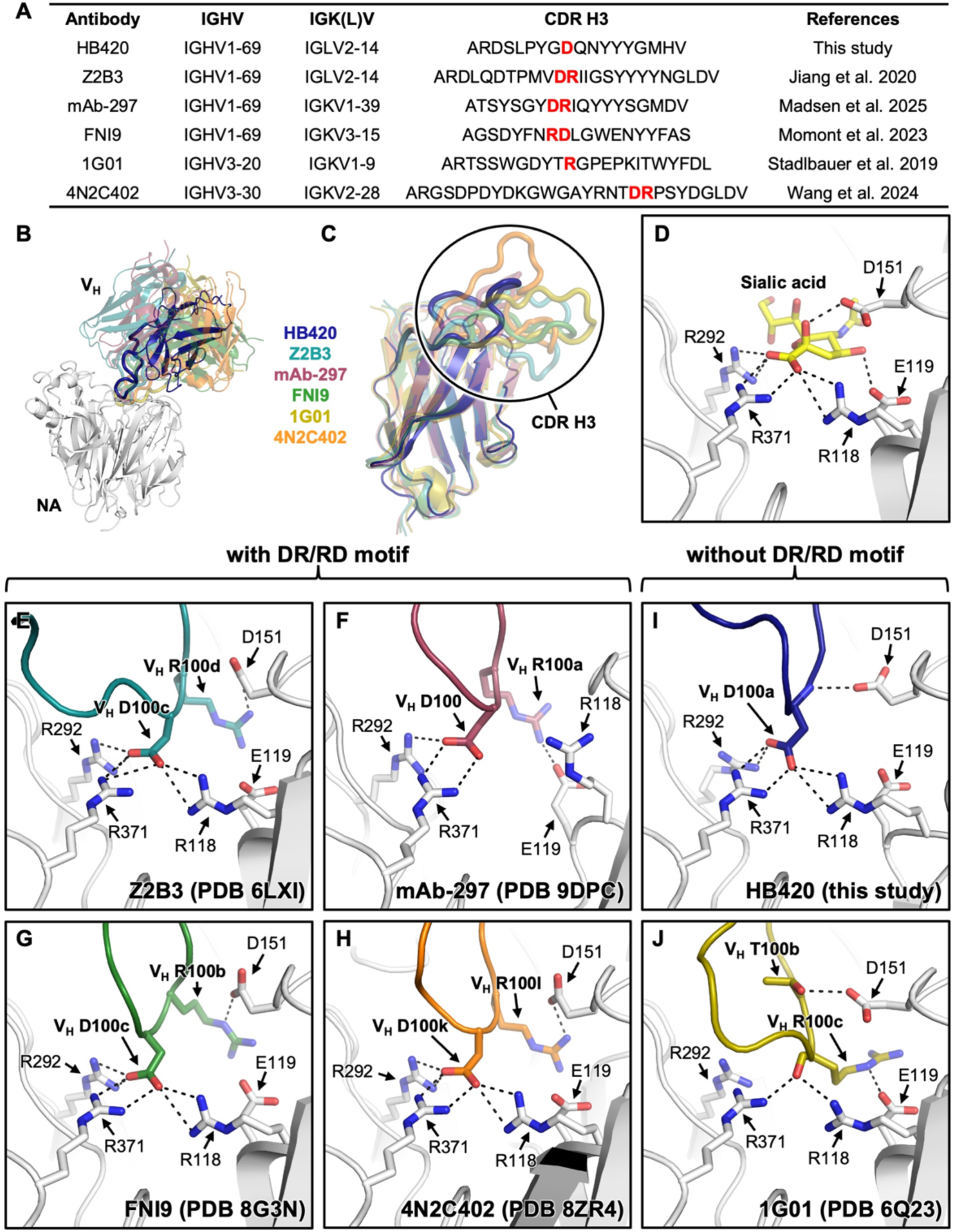
Mimicry of sialic acid binding by HB420. **(A)** Representative cross-group NA antibodies that bind the active site from previous studies.^18,19,24,37,38^ **(B)** Comparison of the angles of approach of the indicated antibodies targeting the NA active site, including HB420 (this study), Z2B3 (PDB 6LXI),^37^ mAb-297 (PDB 9DPC),^24^ FNI9 (PDB 8G3N),^18^ 1G01 (PDB 6Q23),^38^ and 4N2C402 (PDB 8ZR4).^19^ Heavy chain variable domain (V_H_) of each antibody is shown. **(C)** Structural comparison among the indicated antibodies by alignment of the heavy-chain variable domain framework regions. **(D-J)** Interaction of NA active site with **(D)** sialic acid (PDB 8DWB),^68^ **(E)** Z2B3 (PDB 6LXI),^37^ **(F)** mAb-297 (PDB 9DPC),^24^ **(G)** FNI9 (PDB 8G3N),^18^ **(H)** 4N2C402 (PDB 8ZR4),^19^ **(I)** HB420, and **(J)** 1G01 (PDB 6Q23).^38^ H-bonds and electrostatic interactions are represented by black dashed lines.

Recently, several studies have shown that antibodies often utilize a dipeptide motif in the CDR H3, namely aspartic acid-arginine or arginine-aspartic acid (DR/RD), to engage the NA active site.^19,23,24,39^ This DR/RD motif, which is present in Z2B3, mAb-297, FNI9, and 4N2C402, mimic the binding of sialic acid to NA R118, E119, D151, R292, and R371^18,39^ (**Figure 5D**). The Asp in the DR/RD motif electrostatically interacted with NA R118, D151, and R292^18^ (**Figure 5E-H**), whereas the Arg electrostatically interacted with either E119 (**Figure 5F**) or D151 (**Figure 5E, G, H**). While HB420 contained only an Asp (i.e. V_H_ D100a) and not an Arg in its CDR H3, its V_H_ D100a interacted with not only NA R118, D151, and R292, but also NA D151 (**Figure 5I**), which was typically engaged by the Arg in those NA antibodies with the DR/RD motif (**Figure 5E-H**). Same as the Asp in the DR/RD motif of Z2B3, mAb-297, FNI9, and 4N2C402, HB420 V_H_ D100a electrostatically interacted with NA R118, D151, and R292. However, unlike those NA antibodies with the DR/RD motif, the backbone amide group of HB420 V_H_ D100a was positioned close enough to H-bond with NA D151. In other words, the backbone amide group of HB420 V_H_ D100a functioned like the Arg in the DR/RD motif. When we introduced a DR/RD motif into the HB420 by adding an Arg immediately upstream or downstream of V_H_ D100a via insertion or substitution, its binding activity was abolished (**Figure S4**). Together, the interactions formed by a single aspartic acid in the CDR H3 of HB420 were similar to those formed by the DR/RD motif in other NA antibodies.

Besides HB420, 1G01 is another cross-group human antibody that targets the NA active site without a DR/RD motif.^38^ While HB420 encoded an Asp at the tip of the CDR H3, 1G01 had an Arg (V_H_ R100c). The side chain of 1G01 V_H_ R100c electrostatically interacted with NA E119, whereas its backbone carbonyl group H-bonded with R118 and R371 (**Figure 5J**), both of which are typically engaged by the Asp in those NA antibodies with the DR/RD motif. Therefore, the backbone carbonyl group of 1G01 V_H_ R100c functioned like the Asp in the DR/RD motif. Overall, these observations demonstrated that the missing Arg or Asp in the sialic acid mimicry DR/RD motif, as seen in HB420 and 1G01, respectively, could be at least partially compensated by interactions involving the CDR H3 backbone.

### Cross-reactivity of HB420 to H5N1 NA is conferred by heavy chain SHMs

To investigate how HB420 developed cross-reactivity against N1 and N2 subtypes, we evaluated the binding activity of several germline revertants of HB420 (**Figure S5A-B**). HB420 with all SHMs reverted could still bind to Mos99 (H3N2) NA, but showed no detectable binding to Tx24 (H5N1) NA (**Figure 6A and Figure S5C-D**). Additionally, HB420 with only heavy chain SHMs reverted behaved similarly to the fully germline-reverted HB420, whereas HB420 with only light chain SHMs reverted behaved similarly to HB420. We further measured the binding affinity of different germline revertants of HB420 in Fab format against Mos99 (H3N2) NA using biolayer interferometry. Our results showed that HB420 and HB420 with only light chain SHMs reverted had dissociation constants (K_D_s) of less than 1 nM (**Figure 6B and Figure S6**). In comparison, the binding affinities of HB420 with only heavy chain SHMs reverted and HB420 with all SHMs reverted were weaker, with K_D_s of 1.1 nM and 1.8 nM, respectively. Consistent with the binding affinity measurement, HB420 and HB420 with only light chain SHMs reverted showed the strongest virus growth inhibition activity, followed by HB420 with only heavy chain SHMs reverted, and then HB420 with all SHMs reverted (**Figure 6C**). These results demonstrated that SHMs in the heavy chain of HB420 played a critical role in enhancing its binding affinity against H3N2 NA, and concurrently enabled cross-reactivity with H5N1 NA.

**Figure 6.**
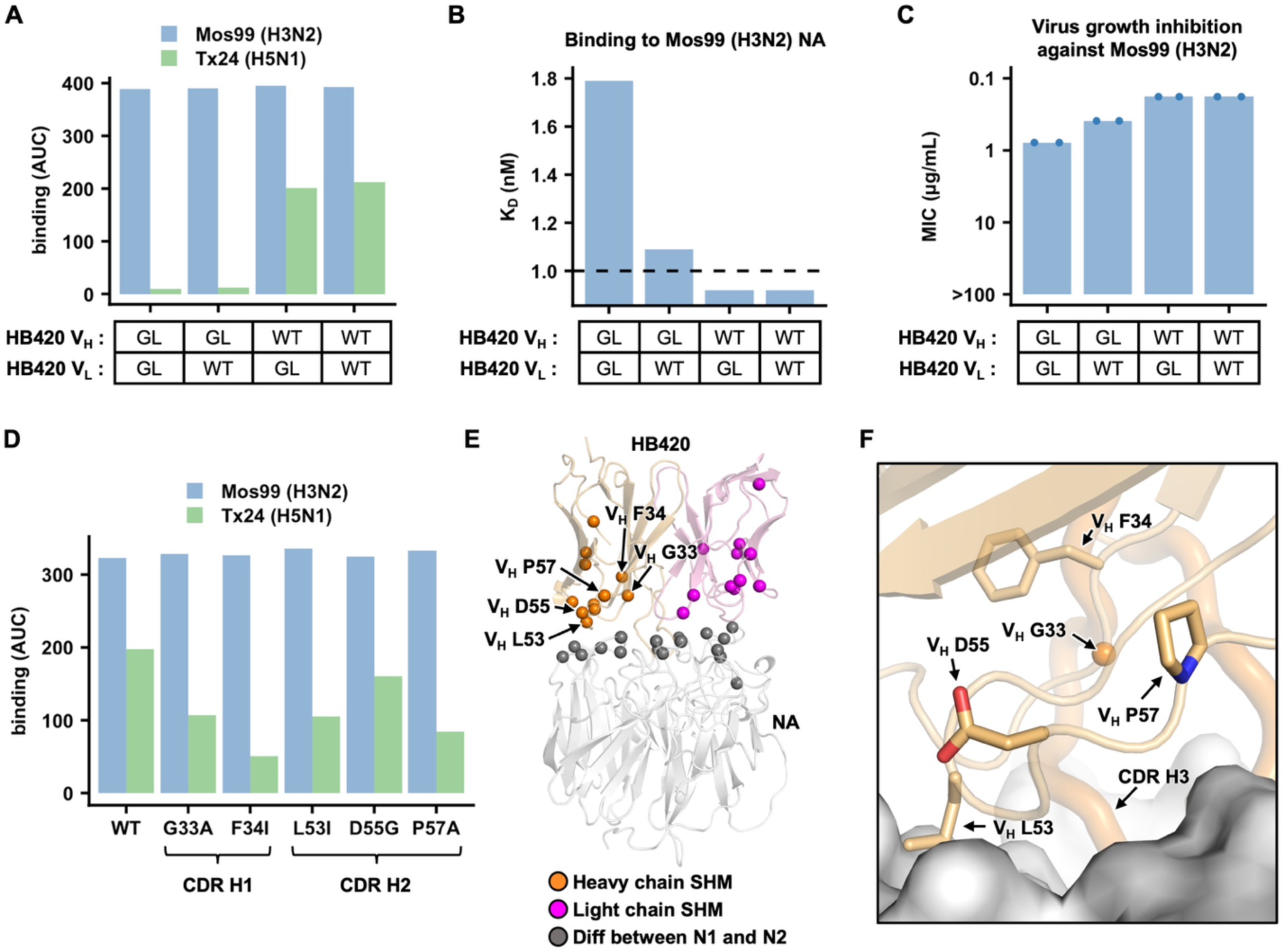
Identification of SHMs that confer cross-reactivity to H5N1 NA. **(A)** Binding of the indicated HB420 germline revertants against recombinant NAs from Mos99 (H3N2) and Tx24 (H5N1) was measured by ELISA. AUC: area under the curve. **(B)** Dissociation constant (K_D_) of the indicated HB420 germline revertants against recombinant Mos99 (H3N2) NA was measured by biolayer interferometry. **(C)** Virus growth inhibition activity of the indicated HB420 germline revertants against Mos99 (H3N2) virus was measured. Minimum inhibitory concentration (MIC) is shown. **(A-C)** WT and GL indicate wild type and germline, respectively. **(D)** Binding of the indicated SHM revertants of HB420 against recombinant NAs from Mos99 (H3N2) and Tx24 (H5N1) was measured by ELISA. **(E)** Spatial locations of SHMs of HB420 are shown. Epitope residues that have different amino acid sequences between Mos99 (H3N2) and Tx24 (H5N1) are shown. **(F)** SHMs of interest on the HB420 heavy chain are indicated on the structure. NA is shown in surface representation. CDR H3 is shown in thick cartoon representation.

To identify SHMs that are important for the cross-reactivity of HB420 with Tx24 (H5N1) NA, we individually reverted the SHMs in its CDRs H1 and H2 to the germline sequence (**Figure S5A**). These SHM revertants include V_H_ G33A and V_H_ F34I in the CDR H1, as well as V_H_ L53I, V_H_ D55G and V_H_ P57A in the CDR H2. While all five of them reduced the binding to Tx24 (H5N1) NA, V_H_ F34I and V_H_ P57A had the strongest effects (**Figure 6D**). Nevertheless, V_H_ residues 34 and 57 were not part of the HB420 paratope (**Figure 6E and Figure S5A**). V_H_ residue 34 were buried, suggesting that SHM V_H_ I34F altered the packing of the protein core, which may in turn change the backbone conformation of the CDRs to enhance binding (**Figure 6F**). The SHM at V_H_ residue 57 replaced an Ala by a Pro (**Figure S5A**), which would rigidify CDR H2 due to the restricted backbone conformational freedom of proline (**Figure 6F**), thereby reducing the entropic cost of binding. These observations showed that non-paratope SHMs played a critical role in conferring HB420 with cross-reactivity to Tx24 (H5N1) NA, despite the sequence variation in the HB420 epitope between the N1 and N2 subtypes (**Figure 6E and Figure S2A**). Notably, such a phenomenon is not uncommon in the evolution of broadly reactive antibodies.^40–43^

## DISCUSSION

Since the first documented human outbreak of HPAI H5N1 virus in Hong Kong in 1997, concerns have persisted about its pandemic potential.^44^ Although sustained human-to-human transmission of HPAI H5N1 virus has not been observed, sporadic cases suggest that limited transmission is possible.^45,46^ The ongoing circulation of HPAI H5N1 clade 2.3.4.4b virus in the US dairy cattle has facilitated its acquisition of multiple mammalian-adaptive mutations.^47^ These developments underscore an unprecedented and urgent pandemic threat posed by HPAI H5N1. According to CDC and WHO, pre-existing population immunity to an emerging influenza virus is one of the key parameters for pandemic risk assessment.^48,49^ While HA-neutralizing antibodies against H5N1 clade 2.3.4.4b are minimal in humans, if not absent, NA-inhibiting antibodies are detected at high level.^27,28^ Studies have shown that these pre-existing H5N1 NA-inhibiting antibodies are likely elicited by H1N1 infection.^27,28,30^ Nevertheless, H5N1 NA-inhibiting antibodies also exhibit a minor, though statistically insignificant, increase following H3N2 infection.^28^ Consistently, although HB420 may not represent a natively paired antibody, it serves as proof-of-concept that NA-inhibiting antibodies can evolve cross-reactivity to H5N1 NA from a germline precursor specific to H3N2 NA. Our data further indicate that the development of such cross-reactivity can arise as a byproduct of affinity maturation against H3N2 NA.

By comparing the characteristics of antibodies targeting the NA active site, we observe many similarities with antibodies that target the HA receptor-binding site. The most notable similarity is their mimicry of sialic acid binding. Several antibodies targeting the NA active site use an aspartic acid residue to mimic the carboxylate group of sialic acid, a strategy also observed in many antibodies targeting the HA receptor-binding site.^50^ Additionally, public clonotype, which is defined as antibodies from different individuals that use the same *IGHV* gene and have similar CDR H3 sequences, seems to be rare in antibodies targeting the NA active site or HA receptor-binding site, in contrast to those targeting the HA stem.^51^ A previous study on antibodies targeting the HA receptor-binding site has shown that they can arise from diverse clonotypes.^52^ The same appears to apply to antibodies targeting the NA active site, based on what is known to date. Finally, since both the HA receptor-binding site and the NA active site are highly conserved among influenza A viruses, antibodies targeting these epitopes can exhibit heterosubtypic cross-reactivity.

Including HB420, the majority of cross-group antibodies targeting the NA active site known to date are encoded by *IGHV1-69*.^18,24,37^ However, the positions of the heavy chain relative to NA vary across these *IGHV1-69* NA antibodies. Additionally, their CDR H3 sequences and conformations are highly diverse. By contrast, although HA stem antibodies also exhibit a bias toward *IGHV1-69*, they often share a similar binding mode by engaging the same hydrophobic pockets in the HA stem through CDR H2 of *IGHV1-69*.^53^ Therefore, it is unclear why *IGHV1-69* is enriched among cross-group NA antibodies targeting the active site. This observation may be due to the small sample size, given the limited number of studies on NA antibodies. It remains to be seen whether this trend holds as additional cross-group NA antibodies targeting the active site are discovered.

Over the past two decades, the discovery and characterization of broadly neutralizing antibodies targeting the HA stem have inspired the development of multiple HA stem-based broadly protective influenza vaccine candidates.^54^ Similarly, knowledge of cross-reactive NA-inhibiting antibodies is important for exploring NA as a target for broadly protective influenza vaccines.^55,56^ An ideal broadly protective influenza vaccine should elicit antibody responses against conserved epitopes on both HA and NA to minimize the potential for resistance. However, our molecular understanding of NA antibodies is way less than that of HA antibodies.^56^ As a result, continued studies of NA antibodies, especially those that confer protection against multiple subtypes, are warranted for the development of broadly protective influenza vaccines.

## ACKNOWLEDGEMENTS

This work was supported by the Carl R. Woese Institute for Genomic Biology Postdoctoral Fellowship (H.L.), the Vallee Scholars Program (N.C.W.), the Searle Scholars Program (N.C.W.), and Howard Hughes Medical Institute Emerging Pathogens Initiative (N.C.W.). We thank Kristen Flatt at the UIUC Materials Research Laboratory Central Research Facilities and Frank Vago at the Purdue Cryo-EM Facility for assistance with cryo-EM experiments.

## AUTHOR CONTRIBUTIONS

H.L., C.K.P.M., and N.C.W. conceived and designed the study. H.L., Y.W.H., Q.W.T., C.C., T.P., A.B.G., M.R.A., J.J.H., R.L., X.C., Y.S., Y.S.T., A.M., M.S., K.J.M., E.X.M., L.E.W., M.T., and L.A.R. performed the experiments. H.L., C.K.P.M., and N.C.W. wrote the paper and all authors reviewed and/or edited the paper.

## METHODS

### Isolation of plasma and peripheral blood mononuclear cells

A total of 245 blood samples from healthy donors were collected through the Red Cross Hong Kong in the year of 2020. Blood samples were first centrifuged at 3000 ×g for 10 min at room temperature to collect plasma samples. The remaining blood was diluted with equal volume of phosphate-buffered saline (PBS), transferred onto the Ficoll-Paque Plus medium (Cytiva), and centrifuged at 400 ×g for 20 min to isolate peripheral blood mononuclear cells (PBMCs), which were subsequently washed with cold RPMI-1640 medium (Thermo Fisher Scientific) three times and stored in cell freezing solution containing 10% dimethyl sulfoxide (DMSO) and 90% fetal bovine serum (FBS) at −80°C until used. All study procedures were performed after informed consent. The study was approved by the Human Research Ethics Committee at The Chinese University of Hong Kong (IRB: 2020.229).

### Construction of the phage display antibody libraries

Two independent phage display antibody libraries (Fab-HLκ and Fab-HLλ) were constructed. In brief, total RNA was extracted from 245 PBMCs using the RNeasy Mini Kit (Qiagen). The cDNAs were then synthesized using the ProtoScript II First Strand cDNA Synthesis Kit (NEB). Using the cDNAs as templates, the heavy chain variable (VH), κ light chain variable (Vκ), and λ light chain variable (Vλ) regions were independently amplified by the AllTaq Master Mix Kit (Qiagen). Subsequently, VH along with the CH1 domain was ligated to Vκ or Vλ along with the CL domain. The ligation products were cloned into a pComb3xTT vector (Addgene plasmid #63891). The phagemid antibody library was electroporated into E. coli TG1 cells (Amid Bioscience), using a Gene Pulser Series II (Biorad) at 1.8 kV, 25 µF and 200 Ω. Transformants were outgrew in 1 mL of SOC medium (Thermo Fisher Scientific) for 1 h at 37°C, and the cells were platted on 2YT agar supplemented with 100 µg mL^-1^ of ampicillin (Thermo Fisher Scientific). The agar plates were incubated overnight at 30°C. Approximately 20 million colonies were collected in 2YT medium supplemented with 30% glycerol. Transformants were diluted in 100 mL of fresh 2YT medium supplemented with 100 µg mL^-1^ ampicillin, giving an OD_600_ of approximately 0.1. Transformants were incubated at 37°C with agitation, until it reached an OD_600_ of approximately 0.45. Hyperphage M13 KO7ΔpIII (Progen, PRHYPE) was added to the transformants at a Multiplicity of Infection (MOI) of approximately 20. The hyperphage-infected transformants were incubated at 37°C for 30 min without shaking, followed by another 30 min at 37°C with shaking at 250 rpm. Kanamycin (Thermo Fisher Scientific) was added at a final concentration of 50 µg mL^-1^, followed by incubation at 27°C for 16-18 h with shaking at 250 rpm. The cultures were centrifuged at 7500 ×g for 10 min at 4°C with a Sorvall LYNX 4000 floor centrifuge (Thermo Fisher Scientific). Supernatant containing the hyperphage was collected and a mixture of polyethylene glycol 6000 and NaCl was added to a final concentration of 4% and 0.5 M respectively, followed by incubation on ice for at least 1 h. A pellet of hyperphage was obtained by centrifuging the mixture at 10,000 ×g for 20 min at 4°C. The pellet was then resuspended in approximately 4 mL of 1× PBS (Corning). Debris was removed by centrifuging the sample at 4000 ×g for 10 min at 4°C. The supernatant was passed through a Millex 0.45 mm syringe filter (Millipore). Precipitation of the hyperphage was repeated as stated above. The final hyperphage pellet, which represented the phage display antibody library, was resuspended in 1× PBS. Quantification of hyperphage was carried out by first treating a serial dilution of phage sample with 0.1 µg mL^-1^ of TPCK-treated trypsin (Thermo Fisher Scientific) for 30 min at 37°C. Then 100 µL of E. coli TG1 outgrew to an OD_600_ of 0.5 was added to the 100 µL of trypsin-treated hyperphage, followed by incubation at 37°C, standing for 30 min and shaking for another 30 min. Serial dilution of the transductants was platted on 2YT agar supplemented with 100 µg mL^-1^ of ampicillin and incubated overnight at 30°C before plaque counting.

### Panning and screening of the phage-display libraries

A total of 100 ng of A/cattle/Texas/56283/2024 (H5N1) NA in 100 µL of 1× PBS was added to each well of an Invitrogen Nunc MaxiSorp flat-bottom 96 well plate (Thermo Fisher Scientific) and incubated overnight at 4°C. After washing off the unbound NA by 1× PBS, each well was filled to the brim with 5% skim milk in 1× PBS and incubated for 2 h at room temperature. The phage display antibody library, either the Fab-HLκ and Fab-HLλ, was diluted to approximately 10^12^ hyperphage mL^-1^ and was incubated with an equal volume of 5% skim milk in 1× PBS for 2 h at room temperature. The phage display antibody library was then added to each well and incubated for 2 h at 37°C. Each well was washed five times with PBST (1× PBS supplemented with 0.1% Tween-20), five times with 1× PBS, and tapped dry on paper towel after each wash. The phage display antibody library was eluted from each well by incubating each well with 0.1 µg/mL of TPCK-treated trypsin for 30 min at 37°C. The eluted phage display antibody library was added to 15 mL of E. coli TG1 culture outgrew to an OD_600_ of 0.45, followed by incubation at 37°C, standing for 30 min and shaking for another 30 min. Transductants were concentrated by centrifuging at 4500 ×g for 10 min, serial diluted, and platted onto 2YT agar supplemented with 100 µg mL^-1^ of ampicillin. Plates were incubated overnight at 30°C. Transductants were collected in 2YT medium supplemented with 30% glycerol and prepared for one more round of screening as described above. The stringency of screening was increased sequentially, by increasing the number of washes to eight times PBST plus seven times 1× PBS for the second round.

### Phage enzyme-linked immunosorbent assay

After the second round of panning, 96 colonies were randomly selected, inoculated into 200 μL of 2YT-GC medium in a 96 deep-well plate and incubated overnight at 30°C, with shaking at 700 rpm using a Incu-MixerTm Microplate vortexer (Benchmark Scientific). Overnight cultures were diluted 1:100 in fresh 2YT medium and incubated at 37°C, with shaking at 700 rpm until the OD_600_ reached approximately 0.5. Subsequently, hyperphage M13 KO7ΔpIII were added to the plates (1:1000 dilution, MOI of around 20) and incubated for 1 h at 37°C. Kanamycin was then added to each well at a final concentration of 50 μg mL^-1^. Cultures were incubated overnight at 28°C, with shaking at 700 rpm. The plates were centrifuged at 2000 ×g. The hyperphage in the supernatant was used for enzyme-linked immunosorbent assay (ELISA). Briefly, a 96-well maxisorp plate (Thermo Fisher Scientific) was coated with 100 ng H5N1 N1 overnight at 4°C. Hyperphage was diluted by equal volume of 5% milk in 1× PBS and incubated in the maxisorp plate at room temperature for 2 h to remove non-specific binders. Similarly, plates coated with the antigen were blocked with 5% skimmed milk in PBST.

After blocking, the plates were incubated with 100 μL of phage supernatant at 37°C for 2 h, before washing three times with PBST. Then 100 μL of anti-M13 gp8 antibody B62-FE2 (catalog #: 61097, Progen) at 1:1000 in PBST was added to each well and the plates were incubated at 37°C for 1 h. Plates were then washed 3× with PBST. Then 100 μL of horseradish peroxidase (HRP)-conjugated anti-mouse IgG antibody (catalog #: 7076S, Cell Signaling Technology) at 1:5000 in PBST was added to each well and plates were incubated at 37°C for 1 h. After the plates were washed five times with PBST, colorimetric detection was performed by adding 100 μL of 1-Step Ultra TMB-ELISA Substrate Solution (Thermo Fisher Scientific) into each well. The reaction was terminated by the addition of 1 M H_2_SO_4_, and the absorbance was measured at 450 nm using a BioTek Synergy HTX Multimode Reader. Clones showing specific binding activity were subjected to Sanger DNA sequencing.

### Cell culture

MDCK-SIAT1 cells (Madin-Darby canine kidney cells with stable expression of human 2,6-sialtransferase, female, Sigma-Aldrich) were cultured in Dulbecco’s modified Eagle’s medium (DMEM) with high glucose (Thermo Fisher Scientific) supplemented with 10% heat-inactivated fetal bovine serum (FBS, Thermo Fisher Scientific), 1% penicillin-streptomycin (Thermo Fisher Scientific), and 1× GlutaMax (Thermo Fisher Scientific). Cell passaging was performed every 3 to 4 days using 0.05% Trypsin-ethylenediaminetetraacetic acid (EDTA) solution (Thermo Fisher Scientific). Expi293F cells (human embryonic kidney cells, female, Thermo Fisher Scientific) were maintained in Expi293 Expression Medium (Thermo Fisher Scientific). Sf9 cells (*Spodoptera frugiperda* ovarian cells, female, ATCC) were maintained in Sf-900 II SFM medium (Thermo Fisher Scientific).

### Expression and purification of NA

The NA head domains, which contained residues 82 to 469 (N2 numbering), were fused to an N-terminal gp67 signal peptide, His6-tag, a vasodilator-stimulated phosphoprotein (VASP) tetramerization domain, and a thrombin cleavage site.^57^ Subsequently, recombinant bacmid DNA was generated using the Bac-to-Bac system (Thermo Fisher Scientific) according to the manufacturer’s instructions. Baculovirus was generated by transfecting the purified bacmid DNA into adherent Sf9 cells using Cellfectin reagent (Thermo Fisher Scientific) according to the manufacturer’s instructions. The baculovirus was further amplified by passaging in adherent Sf9 cells at an MOI of 1. Recombinant protein was expressed by infecting 1 L of suspension Sf9 cells at an MOI of 1. On day 3 post-infection, Sf9 cells were pelleted by centrifugation at 4000 ×g for 25 min, and soluble recombinant NA were purified from the supernatant by affinity chromatography using Ni Sepharose excel resin (Cytiva) and then size exclusion chromatography using a HiLoad 16/100 Superdex 200 prep grade column (Cytiva) in 20 mM Tris-HCl pH 8.0, 100 mM NaCl and 10 mM CaCl_2_. The purified NA protein was concentrated by Amicon spin filter (Millipore Sigma) and filtered by 0.22 µm centrifuge tube filters (Costar). Concentration of the protein was determined by nanodrop (Fisher Scientific). Proteins were subsequent aliquoted, flash frozen by dry-ice ethanol mixture, and stored at −80°C until used.

### Expression and purification of IgG and Fab

The heavy and light chain genes of the obtained antibody were synthesized as eBlocks (Integrated DNA Technologies) and then cloned into human heavy chain either as IgG1 or Fab and human kappa or lambda light chain expression vectors using Gibson assembly according to a previously described method.^58^ The plasmids of heavy chain and light chain were transiently co-transfected into Expi293F cells at a mass ratio of 2:1 (HC:LC) using ExpiFectamine 293 Reagent (Thermo Fisher Scientific). After transfection, the cell culture supernatant was collected at 6 days post-transfection. The IgG and Fab were then purified using a CaptureSelect CH1-XL pre-packed column (Thermo Fisher Scientific).

### Enzyme-linked immunosorbent assay (ELISA)

Nunc MaxiSorp ELISA plates (Thermo Fisher Scientific) were utilized and coated with 100 μL of recombinant proteins at a concentration of 1 μg ml^-1^ in a 1× PBS solution. The coating process was performed overnight at 4°C. On the following day, the ELISA plates were washed three times with 1× PBS supplemented with 0.05% Tween 20, and then blocked using 200 μL of 1× PBS with 5% non-fat milk powder for 2 h at room temperature. After the blocking step, 100 μL of IgGs from the supernatant were added to each well and incubated for 2 h at 37°C. The ELISA plates were washed three times to remove any unbound IgGs. Next, the ELISA plates were incubated with horseradish peroxidase (HRP)-conjugated goat anti-human IgG antibody (1:5000, Invitrogen) for 1 hour at 37°C. Subsequently, the ELISA plates were washed five times using PBS containing 0.05% Tween 20. Then, 100 μL of 1-Step Ultra TMB-ELISA Substrate Solution (Thermo Fisher Scientific) was added to each well. After 15 min incubation, 50 μL of 2 M H_2_SO_4_ solution was added to each well. The absorbance of each well was measured at a wavelength of 450 nm using a BioTek Synergy HTX Multimode Reader.

### Biolayer interferometry binding assay

The binding affinity of Fab to NA protein was measured using an Octet Red96e instrument (Sartorius). The His-tagged NA proteins were loaded onto anti-Penta-HIS (HIS1K) biosensors in kinetics buffer (1× PBS, pH 7.4, 0.01% w/v BSA and 0.002% v/v Tween 20). The binding experiments were performed with the following steps: 1) baseline in kinetics buffer for 60 s; 2) loading of the NAs for 480 s; 3) baseline for 60 s; 4) association of antibody for 120 s; and 5) dissociation of antibody into kinetics buffer for 120 s. Octet assays were carried out at 25°C. Data were analyzed using the Octet Red Data Analysis software version 9.0 and fitted with a 1:1 binding model for the binding data.

### Recombinant virus construction and purification

Using the influenza eight-plasmid reverse genetics system,^59^ recombinant influenza viruses with the NA segment from the indicated strains were generated as 7:1 reassortants on the backbone of A/Puerto Rico/8/1934 (PR8, H1N1). Briefly, DNA plasmids were transfected into a co-culture of HEK293T cells and MDCK-SIAT1 cells (ratio of 6:1) using Lipofectamine 2000 (Thermo Fisher Scientific) and incubated at 37°C for 48 h. Recombinant viruses in the supernatant were plaque-purified on MDCK-SIAT1 cells. Individual plaques were picked and incubated with fresh MDCK-SIAT1 cells, and viral RNAs were extracted from supernatant at 72 h post-infection. The sequences of NA segments were confirmed by Sanger sequencing. All experiments with H5N1 viruses were performed in a biosafety level 3 (BSL3) laboratory at The Chinese University of Hong Kong.

### Enzyme-linked lectin assay (ELLA)

ELLA experiments were performed described.^60^ Briefly, each well of a 96-well microtiter plate (Thermo Fisher Scientific) was coated with 100 μL fetuin (Sigma) at 25 μg mL^-1^ in coating buffer (KPL coating solution; SeraCare) at 4°C overnight. The next day, 50 μL antibodies antibodies at the indicated concentrations in 2-(N-morpholino)ethanesulfonic acid (MES) at pH 6.5, 20 mM CaCl_2_, 1% bovine serum albumin, and 0.5% Tween 20 were mixed with an equal volume of NA protein. This mixture was added to the fetuin-coated plate and incubated for 18 h at 37°C. The plate was then washed six times with PBS with 0.05% Tween 20. Subsequently, 100 μL of HRP-conjugated peanut agglutinin lectin (Sigma-Aldrich) in MES at pH 6.5 with CaCl_2_ and 1% bovine serum albumin was added and incubated for 2 h at room temperature in the dark. The plate was washed six times and developed with 3,3’5,5’-tetramethylbenzidine (TMB) ELISA substrate (Sigma). Absorbance was read at 450 nm using a BioTek Synergy HTX Multimode Reader. Data points were analyzed using Prism software and the 50% inhibition concentration (IC_50_) was defined as the concentration at which 50% of the NA activity was inhibited compared to the negative control.

### Virus growth inhibition assay

MDCK-SIAT1 cells were seeded in a 96-well, flat-bottom cell culture plate (Thermo Fisher Scientific). The next day, serially diluted antibody was mixed with an equal volume of virus and incubated at 37°C for 1 h. The antibody/virus mixture was then incubated with the MDCK-SIAT1 cells at 37°C after the cells were washed twice with PBS. After 1 h incubation, the antibody/virus mixture was replaced with Minimum Essential Medium (MEM) supplemented with 25 mM of 4-(2-hydroxyethyl)-1-piperazineethanesulfonic acid (HEPES), 1 μg mL^-1^ of Tosyl phenylalanyl chloromethyl ketone (TPCK)-trypsin, and antibodies at the same concentration as the initial incubation. The plate was incubated at 37°C for 72 h and the presence of virus was detected by hemagglutination assay. The results were analyzed using Prism software.

### Prophylactic and therapeutic protection experiments

Female BALB/c mice at 6 weeks old (n = 5 mice/group) were anesthetized with isoflurane and intranasally infected with 5× median lethal dose (LD_50_) of recombinant influenza virus with the NA segment from A/Moscow/10/1999 (H3N2) or A/cattle/Texas/56283/2024 (H5N1) on the backbone of PR8 (7:1 reassortant). For determining the protection efficacy against influenza virus with A/Moscow/10/1999 (H3N2) NA, mice were given the HB420 at a dose of 5 mg kg^-1^ intraperitoneally at 4 h before infection (prophylaxis) or 24 h after infection (therapeutic). For determining the protection efficacy against influenza virus with A/cattle/Texas/56283/2024 (H5N1) NA, mice were given the HB420 at a dose of 10 mg kg^-1^ intraperitoneally at 4 h before infection (prophylaxis). Weight loss was monitored daily for 14 days. The humane endpoint was defined as a weight loss of 25% from initial weight on day 0. Of note, while our BALB/c mice were not modified to facilitate interaction with human IgG1, human IgG1 could interact with mouse Fc gamma receptors.^61,62^ To determine the lung viral titers on day 3 post-infection, lungs of infected mice were harvested and homogenized in 1 mL of MEM with 1 μg mL^-1^ of TPCK-trypsin with gentleMACS Dissociator (Miltenyi Biotec). Subsequently, viral titers were measured by TCID_50_ (median tissue culture infectious dose) assay. The results were analyzed using Prism software. The animal experiments were performed in accordance with protocols approved by UIUC Institutional Animal Care and Use Committee (IACUC). H5N1 viral challenge experiment was conducted in The Chinese University of Hong Kong Biosafety Level 3 (BSL3) facility.

### Cryo-EM sample preparation and data collection

The purified NA protein was mixed with the Fab at 1:4 molar ratio (one NA tetramer per four Fabs) and incubated overnight at 4°C before purifying by size exclusion chromatography on the Superose 6 Increased 10/300 column (Cytiva) in 20 mM Tris-HCl pH 8.0, 100 mM NaCl and 10 mM CaCl_2_. The complex peak was concentrated to ∼8 mg mL^-1^. An aliquot of 3 µL sample was applied to a glow-discharged 300-mesh Quantifoil R1.2/1.3 Cu grid and plunge-frozen using a Vitrobot Mark IV (Thermo Fisher Scientific). High-resolution cryo-EM movies were collected on an FEI Titan Krios (Thermo Fisher Scientific) at 300 kV with a K3 detector (Gatan).

### Cryo-EM image processing and model building

Data processing was performed with CryoSPARC^63^ (version 4.5). Movies were subjected to motion correction and CTF estimation, and particles were picked with CryoSPARC blob picker followed by 2D classification. Best classes from blob picker were used as templates for CryoSPARC template pickers, and the resulting particles were cleaned up by multiple rounds of 2D classification before ab initio reconstruction. The best class from ab initio reconstruction was subjected to homogenous refinement, reference-based motion correction, another round of homogenous refinement, local and global CTF estimation, and non-uniform refinement. The map was sharpened with DeepEMhancer,^64^ and all initial atomic model were built using ModelAngelo.^65^ The models were subjected to multiple rounds of manual refinement in Coot^66^ (version 0.9.8) and real-space refinement in Phenix^67^ This process was iterated for several cycles until no significant improvement of the model was observed.

### Data availability

The cryo-EM density map and coordinate of HB420 Fab in complex with Mos99 NA have been deposited to the Electron Microscopy Data Bank (EMDB) with accession code EMD-70808 and the Protein Data Bank (PDB) with accession code 9OSR.

