## Supplementary material for "Evolution of antibody cross-reactivity to influenza H5N1 neuraminidase from an N2-specific germline": SI figure

**Table S1. Cryo-EM data collection and refinement statistics of HB420 Fab in complex with A/Moscow/10/1999 (H3N2) NA.**

| | |
|---|---|
| **Data collection and processing** | |
| Magnification | 81,000 |
| Voltage (kV) | 300 |
| Electron exposure ($e^-$/Å$^2$) | 57.35 |
| Defocus range (μm) | -0.5 to 3.0 |
| Pixel size (Å) | 0.529 |
| Symmetry imposed | C4 |
| Initial particle images (no.) | 547,374 |
| Final particle images (no.) | 355,667 |
| Map resolution (Å)<br>FSC threshold 0.143 | 2.16 |
| **Refinement** | |
| Initial model used (PDB code) | N/A |
| Model composition | |
| Non-hydrogen atoms | 18,806 |
| Protein residues | 2,468 |
| R.m.s. deviations | |
| Bond lengths (Å) | 0.005 |
| Bond angles (°) | 0.622 |
| Validation | |
| MolProbity score | 2.21 |
| Clashscore | 8.52 |
| Poor rotamers (%) | 3.35 |
| Ramachadran plot | |
| Favored (%) | 94.89 |
| Allowed (%) | 5.11 |
| Disallowed (%) | 0.00 |
| **PDB code** | 9OSR |
| **EMDB code** | EMD-70808 |

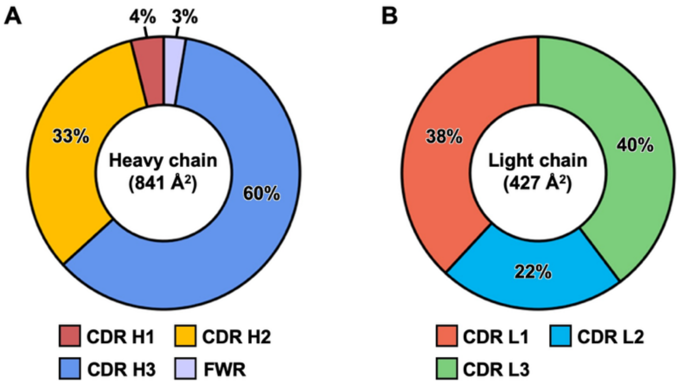


**Figure S1. Composition of the HB420 paratope.** Contributions of different CDRs to the paratope buried surface areas (BSA) of the **(A)** heavy chain and **(B)** light chain of HB420. FWR: framework region.

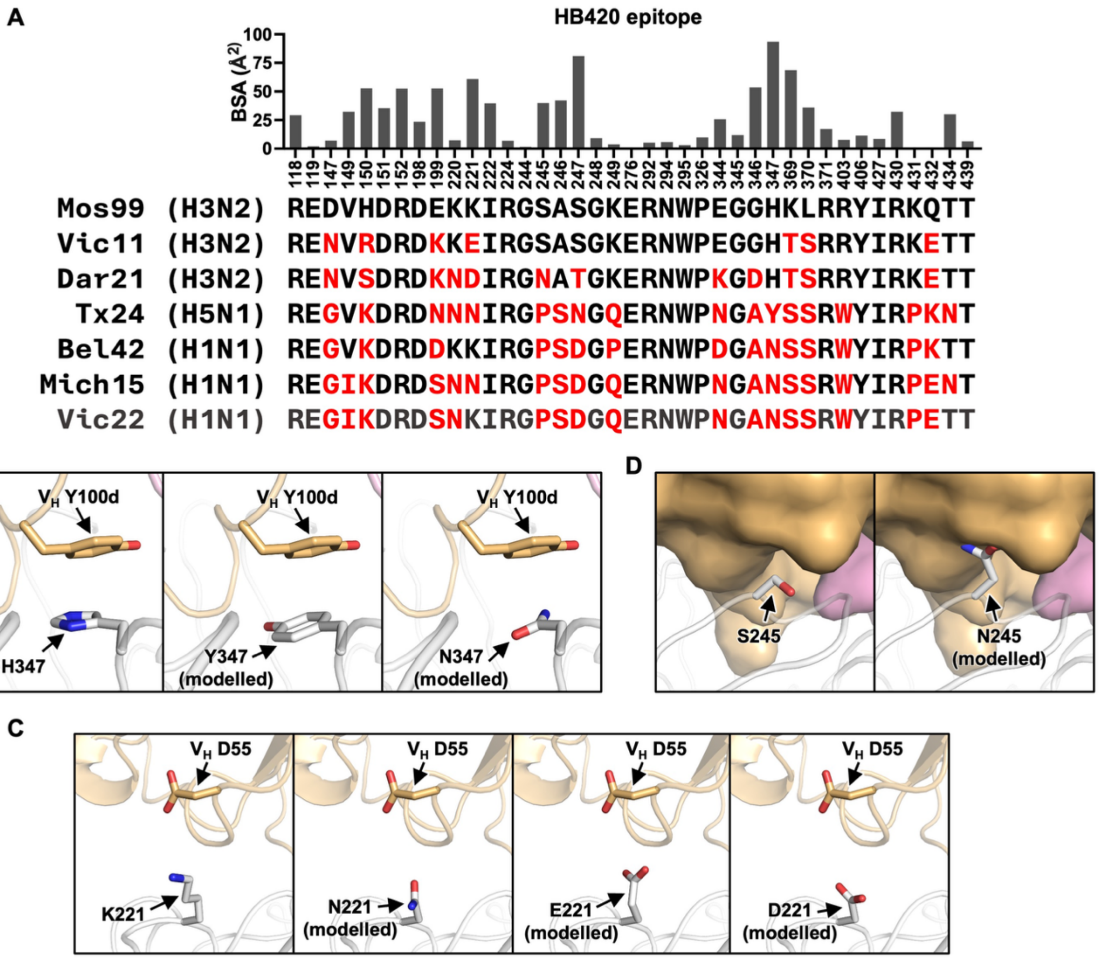


**Figure S2. Sequence variation in the HB420 epitope across influenza strains. (A)** The buried surface area (BSA) of each epitope residue upon binding to HB420 is shown. The amino acid sequence of the indicated strains for each epitope residue is shown at the bottom. Red indicates sequence differences from Mos99 (H3N2). **(B-C)** The structures of HB420 binding to NA with **(B)** Y347 and N347, **(C)** N221, E221, and D221, as well as **(D)** N245 were modelled in PyMOL.

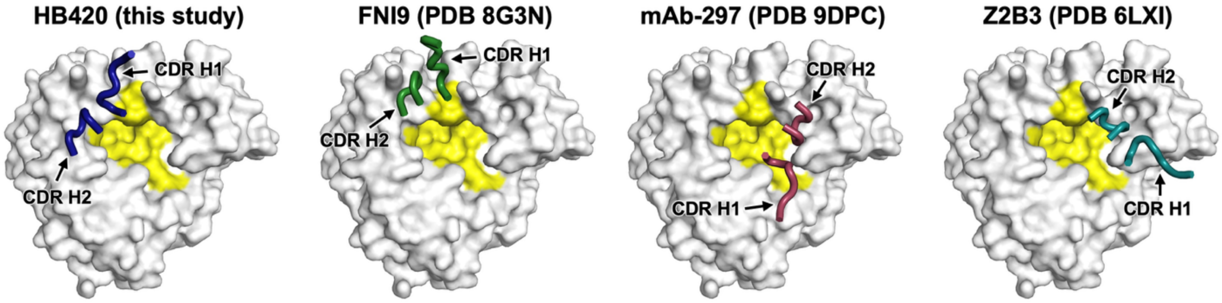


**Figure S3. Positions of CDRs H1 and H2 vary across different antibodies targeting the NA active site.** The positions of CDRs H1 and H2 relative to the NA active site (yellow) are compared across HB420, FNI9 (PDB 8G3N),[1] mAb-297 (PDB 9DPC),[2] and Z2B3 (PDB 6LXI).[3] All structures are shown from the same viewpoint. Only one protomer is shown.

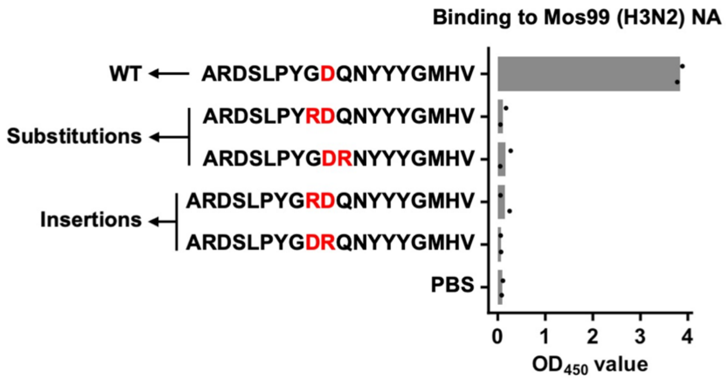


**Figure S4. Introducing a DR/RD motif in the CDR H3 of HB420 abolishes its binding activity.**

The binding activity of different HB420 mutants with a DR/RD motif in the CDR H3 against Mos99 (H3N2) NA is shown. The DR/RD motif was introduced by substitutions or insertions as indicated. The CDR H3 sequence (IMGT numbering) of each mutant is shown. WT: wild type.

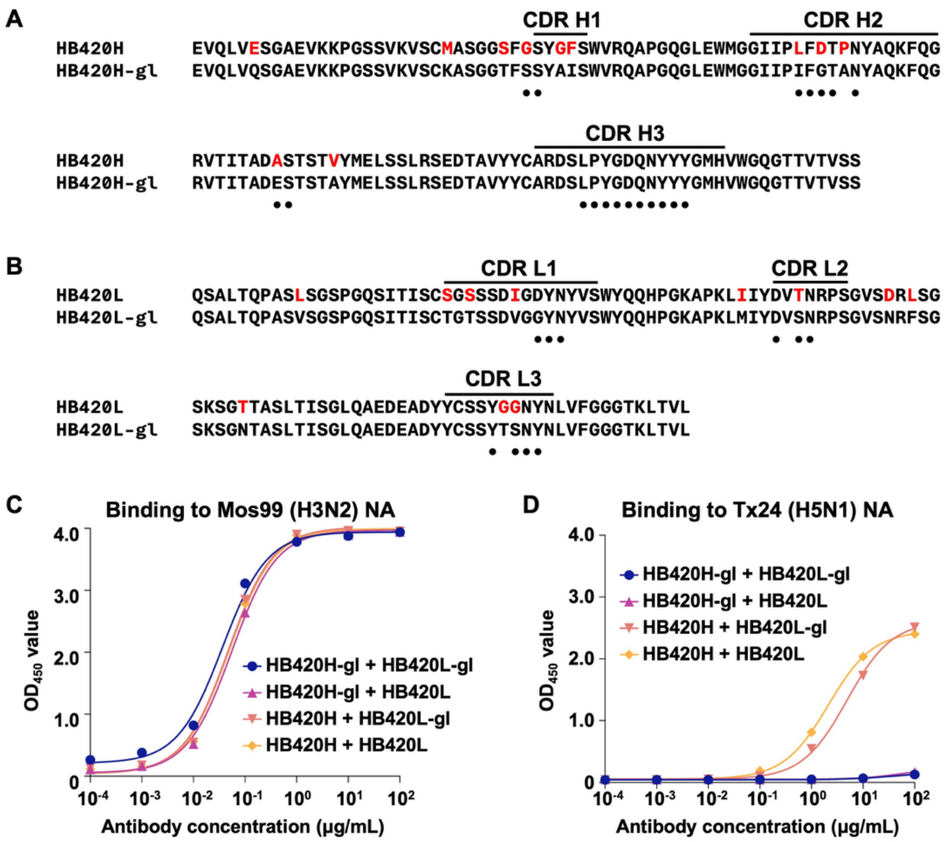


**Figure S5. Binding activity of HB420 germline revertants. (A-B)** Alignment between the amino acid sequence of HB420 and its inferred germline sequence. Residues that represent somatic hypermutations are colored in red. Black dots indicate paratope residues. **(A)** Heavy chain. **(B)** Light chain. **(C-D)** The binding activity of different variants of HB420 in IgG format to **(C)** Mos99 (H3N2) NA and **(D)** Tx24 (H5N1) NA was measured by ELISA. HB420H-gl + HB420L-gl: complete germline revertant (blue). HB420H-gl + HB420L: germline heavy chain with wild-type light chain (purple). HB420H + HB420L-gl: wild-type heavy chain with germline light chain (salmon), HB420H + HB420L: wild type (orange).

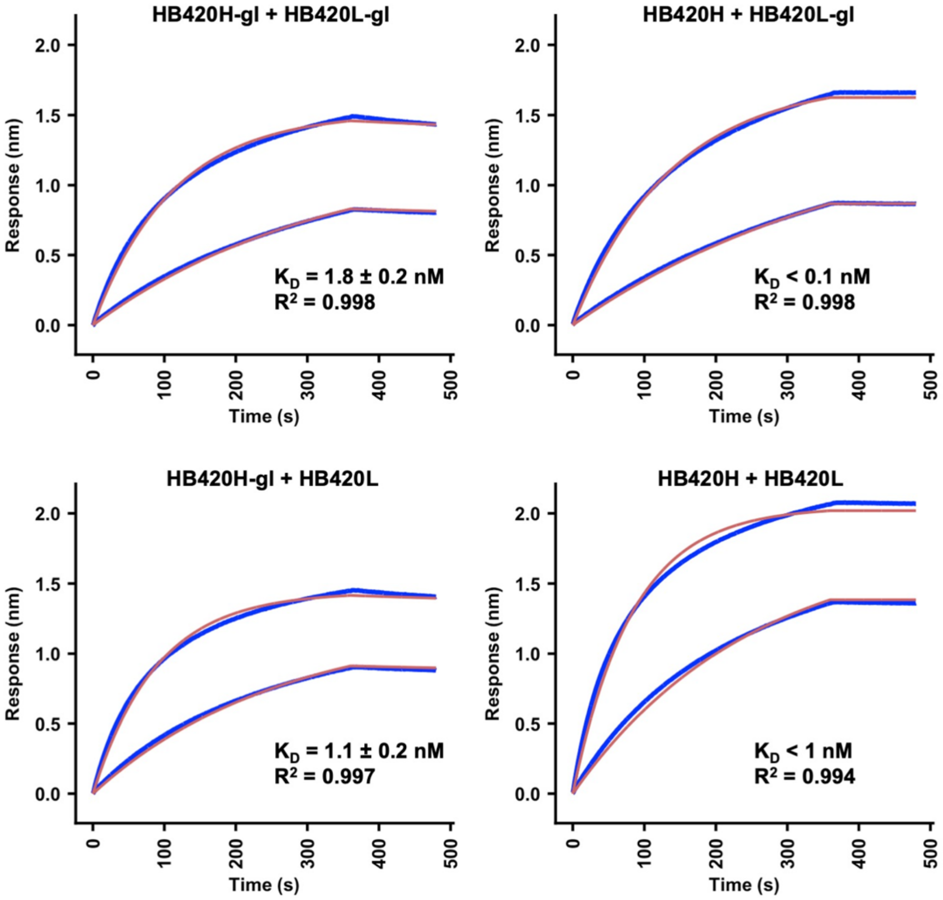


**Figure S6. Sensorgrams for binding of HB420 germline revertants to Mos99 (H3N2) NA.** Binding kinetics of the indicated HB420 germline revertants against Mos99 (H3N2) NA were measured by biolayer interferometry (BLI). Y-axis represents the response. Blue and red lines represent the response curve and the 1:1 binding model, respectively. Binding kinetics were measured for 100 nM and 33 nM of each Fab. Dissociation constants ($K_D$) are shown as mean ± standard deviation. HB420H-gl + HB420L-gl: complete germline revertant. HB420H-gl + HB420L: germline heavy chain with wild-type light chain. HB420H + HB420L-gl: wild-type heavy chain with germline light chain, HB420H + HB420L: wild type.
